# Localized delivery of β-NGF via injectable microrods accelerates endochondral fracture repair

**DOI:** 10.1101/2021.11.16.468864

**Authors:** Kevin O. Rivera, Darnell L. Cuylear, Victoria Duke, Kelsey Marie O’Hara, Bhushan N. Kharbikar, Alex N. Kryger, Theodore Miclau, Chelsea S. Bahney, Tejal A. Desai

## Abstract

Currently, there are no biological approaches to accelerate bone fracture repair. Osteobiologics that promote endochondral ossification are an exciting alternative to surgically implanted bone grafts, however, the translation of osteobiologics remains elusive because of the need for localized and sustained delivery that is both safe and effective. In this regard, an injectable system composed of hydrogel-based microparticles designed to release osteobiologics in a controlled and localized manner is ideal in the context of bone fracture repair. Here, we describe poly (ethylene glycol) dimethacrylate (PEGDMA)-based microparticles, in the form of microrods, engineered to be loaded with beta nerve growth factor (β-NGF) for use in a murine tibial fracture model. In-vitro studies demonstrated that protein-loading efficiency is readily altered by varying PEGDMA macromer concentration and that β-NGF loaded onto PEGDMA microrods exhibited sustained release over a period of 7 days. In-vitro bioactivity of β-NGF was confirmed using a tyrosine receptor kinase A (Trk-A) expressing cell line, TF-1. Moreover, TF-1 cell proliferation significantly increased when incubated with β-NGF loaded PEGDMA microrods versus β-NGF in media. In-vivo studies show that PEGDMA microrods injected into the fracture calluses of mice remained in the callus for over 7 days. Importantly, a single injection of β-NGF-loaded PEGDMA microrods resulted in significantly improved fracture healing as indicated by significant increases in bone volume, trabecular connective density, and bone mineral density and a significant decrease in cartilage despite a remarkably lower dose (∼111 fold) than the β-NGF in media. In conclusion, we demonstrate a novel and translational method of delivering β-NGF via injectable PEGDMA microrods to improve bone fracture repair.

## Introduction

Bone fractures are one of the most common injuries worldwide, affecting millions of individuals, and imparting billions of dollars in healthcare cost burden annually^1,2^. Although bones possess the ability to fully regenerate, fractures induce significant pain and loss in quality of life throughout the time course of healing, which can take 3-6 months under normal conditions^3^. Delayed healing or failure to heal (non-unions) occur in 5-10 % of cases leading to healing times often exceeding 9 months and/or requiring multiple surgeries to achieve union^4^. Co-morbidities such as age, smoking, diabetes, and obesity drastically increase the prevalence of impaired bone healing to upward of 50 % of fracture patients^5,6^. Currently, surgical application of autologous bone grafts or adjustments to the orthopaedic hardware are the standard of care approaches to stimulate healing. Surgical intervention is associated with significant challenges, including, donor site morbidity, longer recovery time, increased costs, and risk of infection, among others^7,8^. There are no pharmacologic reagents approved to accelerate fracture healing or prevent non-union^1^. Given these limitations with surgical methods, there is an unmet clinical need to develop pharmacological approaches to improve bone regeneration without surgery.

Bone morphogenetic proteins (BMPs) are perhaps the most established osteoanabolic growth factors and have been used for regenerating bone through a direct effect on osteoblastic differentiation of osteochondral progenitors^9^. Of the BMPs, BMP-2 is the only osteobiologic growth factor with FDA approval for treatment of problematic fractures^10^. Currently clinical application of BMP-2 in fracture repair remains limited due to its uncertain efficacy, high cost, and growing list of adverse outcomes that includes inflammatory complications, ectopic bone formation, and osteolysis^11,12^. The serious complications are believed to be driven by the supraphysiological doses needed to achieve therapeutic efficacy and lack of drug delivery approaches to mitigate side effects. Thus, it is imperative to explore the use of novel osteobiologics to accelerate fracture repair and to develop optimal drug delivery systems to localize the biologic’s effects and mitigate severe adverse outcomes.

While therapeutic use of BMPs have focused on promoting direct bone formation (intramembranous ossification), bone fractures heal primarily through endochondral ossification (EO). EO, or indirect bone formation, occurs when an avascular, aneural cartilage intermediate converts into a vascularized and innervated bone. EO fracture repair is a dynamic process that proceeds through four overlapping steps: 1) formation of hematoma at the fracture site and activation of the proinflammatory phase, 2) formation of a fibrocartilaginous callus via chondrocytes, 3) formation of a bony callus via chondrocyte hypertrophy and mineralization, and 4) remodeling and compact bone formation^13–16^. Although the molecular mechanisms governing chondrogenesis and hypertrophy are well described, it remains unclear which mechanisms regulate the conversion of cartilage to bone^17,18^. Many molecular pathways have been identified to regulate cartilage to bone transformation including transforming growth factor β (TGF-β), Sox9, Runx2, Wnt, and Indian hedgehog (Ihh)^19–30^.

Recently, nerve growth factor (NGF) signaling via its high affinity receptor tropomyosin receptor kinase A (TrkA) has been shown to be important in both intramembranous and endochondral ossification. In stress fractures, which heal through intramembranous ossification, NGF/TrkA signaling triggers reinnervation, vascularization, and osteoblastic activity during repair^31^. Moreover, NGF/TrkA signaling is now known to be crucial in long bone development as well as skeletal adaptations to mechanical loads^32,33^. Together these studies suggest that NGF could be used as an ostebiologic drug to promote long bone fracture repair by synergistically promoting intramembranous and endochondral bone repair. Towards this, our group has detailed the temporal expression of NGF and TrkA during long bone fracture repair and identified the cartilaginous phase as a key timepoint for local exogenous injections of recombinant beta NGF (β-NGF) to accelerate endochondral fracture repair^34^. In this system, β-NGF delivery promoted cartilage to bone transformation by upregulating the Wnt and Ihh pathways^34^. However, as our published work demonstrated, drug administration by local injection required high dosages and repeated administration to elicit therapeutic effects^34^. Such dosing regimens will likely reduce patient compliance and creates an opportunity to develop controlled delivery platforms that can achieve sustained drug delivery in the absence of surgical implantation^35,36^.

Localized drug delivery to the fracture site with the use of injectable microparticles made of polymeric hydrogels is a highly translatable strategy to overcome the limitations associated with conventional drug administration. Poly (ethylene glycol) (PEG)-based hydrogels are clinically relevant polymer systems based on their low-immunogenic, non-cytotoxic, and biocompatible properties^37^. Therefore, PEG has had extensive biomedical uses for years, particularly drug delivery. Modifying PEG hydrogels with methacrylation (PEGDMA) enables these monomers to be easily formed into three-dimensional (3D) polymers with defined shapes and size through initiation of free radical chain photopolymerization of the methacrylate groups at each end of the PEG chain^38^. Proteins can be readily loaded into PEGDMA microparticles to allow for localized and sustained release, based on the crosslinking density of the polymer^39^. Previously, our group has fabricated high aspect ratio PEGDMA microrods by photolithography for targeted drug delivery^40,41^. High-aspect ratio particles, such as microrods, are an exciting class of microparticles that provide a high surface area to volume ratio and form localized 3D porous networks upon injection, which can maximize local cellular interactions and provide open porous systems for localized drug delivery, cellular recruitment, and activation^42^. Additionally, high-aspect ratio microparticles evade internalization by phagocytic cells such as macrophages^43,44^.

In this work, we build on our foundational research showing recombinant β-NGF promotes endochondral repair in a murine fracture model^34^ by encapsulating the protein in the PEGDMA microrods for sustained local delivery. We hypothesized that local and sustained release of β-NGF from PEGDMA microrods promotes enhanced endochondral fracture repair relative to soluble β-NGF and untreated controls. To test this hypothesis, we first aimed to define key characteristics of the PEGDMA microrod polymer mesh network density for maximization of β-NGF loading efficiencies. By varying concentration of the PEGDMA precursor solution to produce unique network densities, we found that higher % PEGDMA (v/v) microrods had increased loading efficiencies and more sustained release of β-NGF over 7 days when compared to lower % PEGDMA (v/v) microrods. To confirm β-NGF encapsulated within these microrods maintains bioactivity upon release we utilized *in vitro* studies with erythroleukemia cells, a TrkA expressing cell line, to demonstrate retained cellular proliferation relative to other treatment groups. Subsequently, we established these PEGDMA microrods can be delivered and retained within or adjacent to the fracture callus for at least 7 days post-injection. Efficacy studies using our established murine model demonstrate that the mice receiving β-NGF loaded PEGDMA microrods exhibited improved fracture repair relative to soluble β-NGF or negative control. Herein, we describe an injectable system for localized β-NGF delivery for accelerating endochondral fracture repair.

## Methods

### PEGDMA microrod fabrication

Microrods were fabricated as previously described^45^. Briefly, poly (ethylene glycol) dimethacrylate (PEGDMA molecular weight 750, photoinitiator 2,2-dimethoxy-2-phenylacetophenone (DMPA) were dissolved in 1-vinyl-n-pyrrolidone (NVP) to a concentration of 100 mg/mL in phosphate-buffered saline (PBS). 25, 75, and 90 % PEGDMA microrods were created by varying the % v/v PEGDMA to PBS. Photolithography was used to create microrods designed to have the dimensions 100×15×15 μm, micro-fabricated on 3-inch silicon wafers. The PEGDMA precursor solution was deposited onto each wafer wherein the wafers had a 15 μm-deep bevel prefabricated with SU-8 2015. The wafer was exposed using a Karl Suss MJB3 mask aligner to a 405 nm light source through a microrod patterned photomask at 9 mW/cm^2^. Microrods on the wafer were rinsed and removed with 70 % ethanol while gently scraping with a cell scraper. The collected microrods were centrifuged and rinsed in 70 % ethanol three times before being resuspended in sterile deionized water (diH_2_O) with 10 % sucrose and 0.05 % tween-20 to prevent aggregation. Aliquots of ∼100,000 PEGDMA microrods were then lyophilized, sealed, and stored at 4 °C until further use. A subset of PEGDMA microrods was resuspended in PBS and micrographed under bright field (BF). Another subset was stained with the low molecular weight dye 4′,6-diamidino-2-phenylindole (DAPI) 1 μg/mL in PBS for 5 minutes, washed with PBS gently three times, then immediately imaged using a Nikon Ti microscope.

### Protein loading of PEGDMA microrods

Bovine α-Chymotrypsinogen A (Sigma-Aldrich) was used as a proxy for comparing loading efficiencies between 25, 75, and 90 % PEGDMA microrods as its molecular weight (25.7 kDa) is similar to that of β-nerve growth factor (β-NGF, 27 kDa). Lyophilized PEGDMA microrods aliquots (∼100,000 microrods/aliquot) were resuspended in 20 μL of 1 mg/mL α-Chymotrypsinogen A in diH_2_O. After resuspension, the microrods were allowed to passively adsorb protein for 30 hours in 4 °C. After loading, microrods were resuspended in diH_2_O, gently spun down to pellet in tube, and the supernatants were used to perform a micro bicinchoninic acid (μBCA) protein assay to quantify the amount of protein left in solution. To determine loading efficiency, the following equation was used:

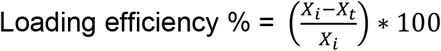

Where X_i_ is the quantity of drug added initially during preparation and X_t_ is amount of protein in the supernatant after 30 hours. Loading of PEGDMA microrods with β-NGF and calculation of loading efficiency was done similarly in subsequent assays.

### Erythroblast (TF-1) Cell Proliferation

TF-1 cell proliferation assay was modified from the established method^46^. Briefly, TF-1 cells (ATCC) were cultured for 7 days in RPMI 1640 Medium modified with 2 mM L-glutamine, 10 mM HEPES, 1 mM sodium pyruvate, 4,500 mg/L glucose, and 1,500 mg/L sodium bicarbonate (ATCC) supplemented with 2 ng/ml recombinant human Granulocyte-Macrophage Colony-Stimulating Factor (Sigma-Aldrich) and 10 % fetal bovine serum. Following 7 days of cell growth, confluent TF-1 cells were collected by centrifugation and resuspended into 24-well microplates with 30,000 cells per well containing 600 μl of serum-free medium. Cells were cultured in serum-free medium for 24 hours to synchronize the cells prior to adding treatment groups. After 24 hours, high pore density (0.4 micron) transwell inserts containing either 100 μl of serum-free medium, 16,000 empty microrods, 2,000 ng of soluble β-NGF, or 16,000 microrods containing 18 ng of β-NGF were inserted into each well and cultured in these conditions for an additional 96 hours. A 24-well plate containing 30,000 cells per well was removed and analyzed (see below) after the 24 hour serum-free medium incubation as a Day 0 control. After 96 hours, the transwell inserts were removed and 300 μl from each well was aspirated. The 300 μl collected was aliquoted into an individual well in a 96-well microplate containing 100 μl each (3 wells total for a single well in the 24-well plate) and subjected to a CyQuant© Direct Proliferation Assay Kit (Thermo Fisher) per the manufacturers protocol. Data is represented as a fold change relative to the cell number at Day 0.

### β-NGF elution from PEGDMA microrods

Lyophilized 90 % PEGDMA microrods were loaded by resuspending ∼100,000 (100 k) microrod aliquots in 20 μL of 1 mg/mL recombinant human β-NGF (Peprotech) as described in the *Protein loading of PEGDMA microrods* section. After loading, microrods were gently rinsed thrice with PBS. PEGDMA microrods were then further divided, samples consisted of 16k microrods/microtube suspended in 250 μL of PBS (pH 7.4). Samples placed onto an orbital shaker (100 RPM) within an incubator (37 °C). PEGDMA microrods were spun down gently and the supernatants were collected and replenished at 6, 24, 48, 72, 96, 120, 144, and 168 hours. Collected supernatants were immediately flash frozen and stored at −80 °C until further use. ELISAs for human β-NGF (RayBiotech) were performed per the manufacturer’s instructions and daily release amounts were calculated by an established standard curve.

### Murine tibial fracture model

Approval was obtained from the University of California, San Francisco (UCSF) Institutional Animal Care and Use Committee (IACUC) prior to performing the mouse studies and the procedures were carried out in accordance with approved guidelines and regulations. Studies were conducted on the C57BL6/J wild type strain obtained from Jackson Labs. Briefly, adult (10-16 weeks) male mice were anesthetized via inhalant isoflurane, and closed non-stable fractures were made mid-diaphysis of the tibia via three-point bending fracture device^47^. Fractures were not stabilized as this method promotes robust endochondral repair. After fractures are created, animals were provided with post-operative analgesics (buprenorphine sustained release). Animals were socially housed and allowed to ambulate freely.

### Local injections

Percutaneous injections into tibial fracture calluses of mice were administered 7 days post-fracture. A precise microliter Hamilton© syringe was utilized for all injections wherein experimental agents were injected in 20 μL of PBS. Experimental groups are as follows: Controls injected with sterile PBS, β-NGF group injected with 500 ng of β-NGF in PBS, non-loaded microrods group injected with 16,000 PEGDMA microrods in PBS, and β-NGF microrods group injected with 16,000 PEGDMA microrods loaded with 18 ng of β-NGF.

### Micro-Computed tomography (μCT)

μCT analysis was performed as previously described^48,49^. Fracture tibias were dissected free of attached muscle 14 days post-fracture, fixed in 4 % PFA and stored in 70 % ethanol. Fracture calluses were analyzed in the UCSF Core Center for Musculoskeletal Biology (CCMBM, NIH P30 funded core) using the Scanco μCT50 scanner (Scanco Medical AG, Basserdorf, Switzerland) with 10 μm voxel size and X-ray energies of 55 kVp and 109 μA. A lower excluding threshold of 400 mg hydroxyapatite (HA)/mm^3^ was applied to segment total mineralized bone matrix from soft tissue in studies of control and treated mice. Linear attenuation was calibrated using a Scanco hydroxyapatite phantom. The regions of interest (ROI) included the entire callus without existing cortical clearly distinguished by its anatomical location and much higher mineral density. μCT reconstruction and quantitative analyses were performed to obtain the following structural parameters: volume fraction (bone volume/total volume as %), trabecular connective density as trabecular bifurcations (#/mm^3^), and bone mineral density (mg HA/cm^3^).

### Localization histology

Tibias were harvested 12,14, and 21 days post-fracture (5, 7, or 14 days post-injection of microrods) to observe microrod localization. At time of collection, tibias were fixed in 4 % PFA and decalcified in 19 % EDTA for 14 days at 4 °C with rocking and solution changes every other day. Tibia were processed for paraffin embedding, serial sections were cut at 10 μm (3 sections per slide), and Hall-Brundt’s Quadruple (HBQ) staining protocol was done to visualize the bone (red) and cartilage (blue) as previously described to localize PEGDMA microrods^34,49^. The sections were mounted on slides with Permount™ mounting medium and brightfield images were captured on a Leica DMRB microscope.

### Histomorphometry

Fracture callus composition was determined using quantitative histomorphometry of tibia harvested 14 days post-fracture. Standard principles of histomorphometric analysis^49^ were utilized to quantify the bone and cartilage fraction in the fracture callus using the first section from every 10^th^ slide analyzed, such that sections were 300 μm apart. Images were captured using a Nikon Eclipse Ni-U microscope with Nikon NIS Basic Research Elements Software version 4.30. Quantification of callus composition (cartilage, bone, fibrous tissue, background) was determined using the Trainable Weka Segmentation add-on in Fiji ImageJ (version 1.51.23; NIH, Maryland, USA)^50^. Volume of specific tissue types was determined in reference to the entire fracture callus by summing the individual compositions relative to the whole.

### Statistical analysis

Individual dots on graphs represent biological replicates, error bars represent standard error of the mean (SEM). Measurements were taken from distinct samples. Data were analyzed using GraphPad Prism (version 8, GraphPad Software, San Diego, CA). ANOVA was used to determine statistical differences between multiple groups followed by Tukey’s HSD post-hoc comparison testing. Significant differences were defined at p < 0.05.

## Results

### Injectable PEGDMA microrod fabrication via photolithography

PEGDMA microrods were fabricated through a process of photolithography (**Fig. 1**). The exact dimensions of the PEGDMA microrods can be carefully controlled by the photomask (length and width). The microrod height is determined by distance between the silicon wafer and the photomask which is manually controlled by use of the mask aligner. PEGDMA microrods are then cross-linked with UV irradiation, detached with a cell scraper from the silicon wafer and collected. Each individual PEGDMA microrod had the following dimensions: *H =* 15 μm, *W* = 15 μm, and *L* = 100 μm. The 3D structure of the microrods is formed through free radical chain photopolymerization of the methacrylate groups at each end of each 750 MW PEG monomer unit. This system provides a high-throughput method to produce highly uniform PEGDMA microrods.

**Figure 1.**
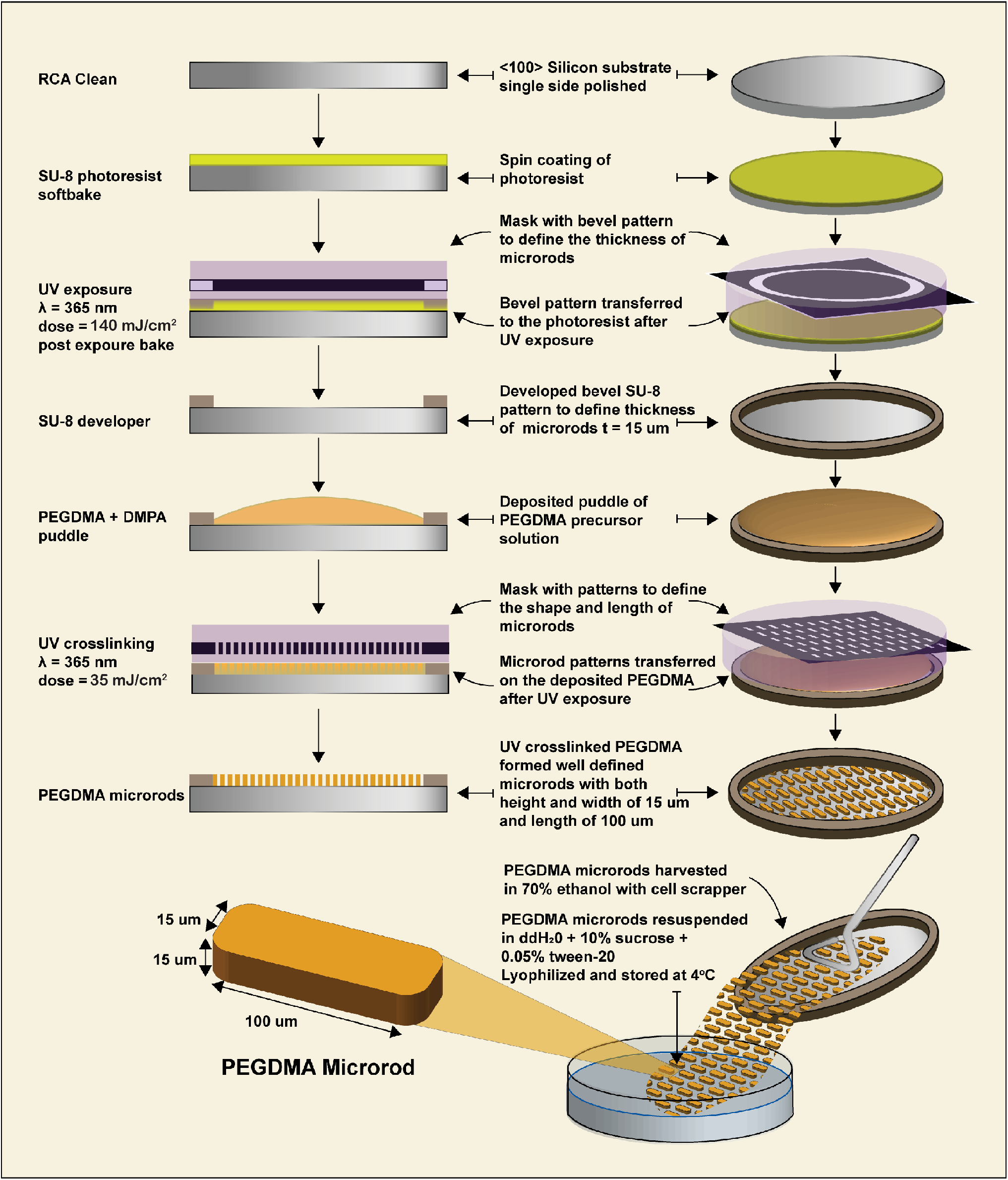
Schematic of PEGDMA microrod fabrication via photolithography.

### PEGDMA microrod macromer concentration changes protein-loading efficiency

The first goal was to tune the PEGDMA microrod polymer network density to maximize protein loading efficiency. Prior to photolithography, the PEGDMA macromer concentration was adjusted to contain low (25 %), medium (75 %), or high (90 %) volume (v/v) amounts (**Fig. 2A**). After fabrication, the PEGDMA microrods were lyophilized (freeze dried) to remove any residual liquid remaining that could alter protein loading and then loaded with α-Chymotrypsinogen A as a proxy protein, as its molecular weight is similar to that of β-NGF (26 kDa). The low and medium macromer volume PEGDMA microrods had modest loading efficiencies of less than 5 % and 15 %, respectively. The high macromer volume PEGDMA microrods exhibited a significantly larger loading efficiency, over 30 %, when compared to the other PEGDMA concentrations. Since the 90 % PEGDMA (v/v) microrods had the best loading efficiency, we chose this formulation for all the subsequent experiments. We then confirmed β-NGF loading efficiency by loading high macromer PEGDMA microrods with β-NGF which resulted in over 40 % loading efficiency (Supplemental Fig. 1). DAPI, a commonly used nuclear counter stain, can be easily adsorbed into the PEGDMA microrods and visualized with fluorescent microscopy (**Fig. 2B**). The DAPI-stained microrods are uniform in size and do not exhibit any aggregation, indicating good dispersity in solutions to allow for more optimal protein loading. Given that protein loading is driven by physisorption, we wanted to qualitatively examine the absorption of proteins onto the microrods and the rate at which protein elution occurs after loading. Using FITC-BSA as a model protein, the superficial layer of the PEGDMA microrods are coated with the fluorescently tagged protein with no diffusion at time 0 (**Fig. 2C, Top)**. After 60 minutes of incubation, diffusion is drastically increased and FITC-BSA can be observed eluting in the surrounding space of the PEGDMA microrods (**Fig. 2C, Bottom**).

**Figure 2.**
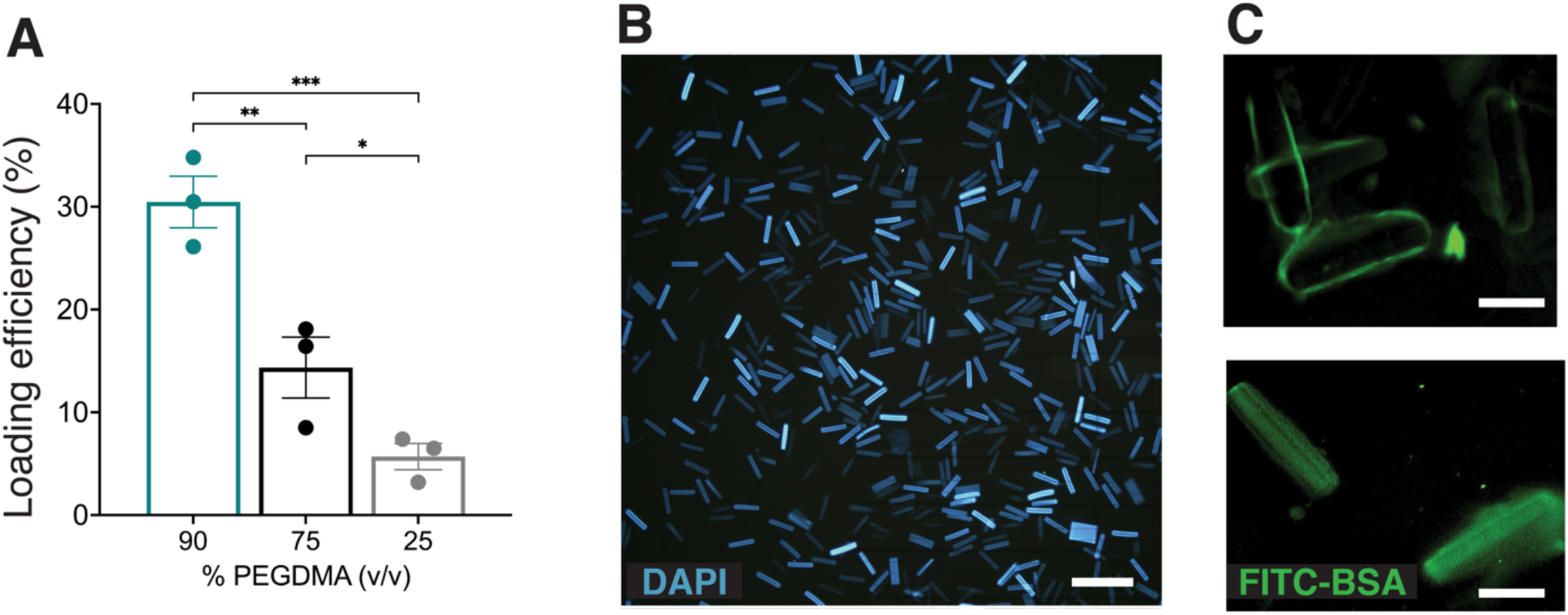
Lyophilized PEGDMA microrods are readily protein loaded. (A) Loading efficiency significantly increases with increasing concentration of PEGDMA (% PEGDMA v/v). Data shown as means with error bars representing SEM. *p<0.05, **p<0.01, ***p<0.001 determined by ANOVA with Tukey’s post hoc test for multiple comparisons (n=3) (B) DAPI-loaded 90% PEGDMA microrods, scale bar = 25μm. (C) Fluorescent micrographs of FITC-BSA loaded in 90% PEGDMA microrods taken after 0 mins (top) and after 60 mins of incubation at room temperature (bottom), scale bars = 50μm.

### Bioactivity retention and sustained release of β-NGF from PEGMDA microrods

We next tested whether β-NGF retained bioactivity when released from the PEGDMA microrods. To do this, we utilized the erythroleukemia TrkA expressing cell line, TF-1, in the presence of culture media (control), 2,000 ng of soluble β-NGF, non-loaded microrods, or 16,000 PEGDMA microrods loaded with 18 ng of β-NGF (**Fig. 3A**). We hypothesized that sustained release from β-NGF microrods would increase proliferation of TF-1 cells relative to the other treatment groups following 4 days of culture. As expected, soluble β-NGF treated cells exhibited a 2-fold increase in proliferation relative to the control. Interestingly, the non-loaded microrods also exhibited a similar increase in proliferation as soluble β-NGF treated cells. However, our hypothesis was supported by the statistically significant 4-fold increase in proliferation by the β-NGF microrods. The increase in proliferation may be attributed to the sustained release of β-NGF which we observed over a 168 hour (7 day) period (**Fig. 3B & C**). The β-NGF microrods exhibited an initial burst release of β-NGF within the first 24-48 hours, followed by sustained release over the next 120 hours (days 2-7), as measured by ELISAs (**Fig. 3B**). The total daily amount of eluted β-NGF decreased with time indicating concentration dependent (first order) release kinetics from the PEGDMA microrods. Nonetheless, we were able to detect and quantify elution of β-NGF over a 7 day period (168 hours) (**Fig. 3C**).

**Figure 3.**
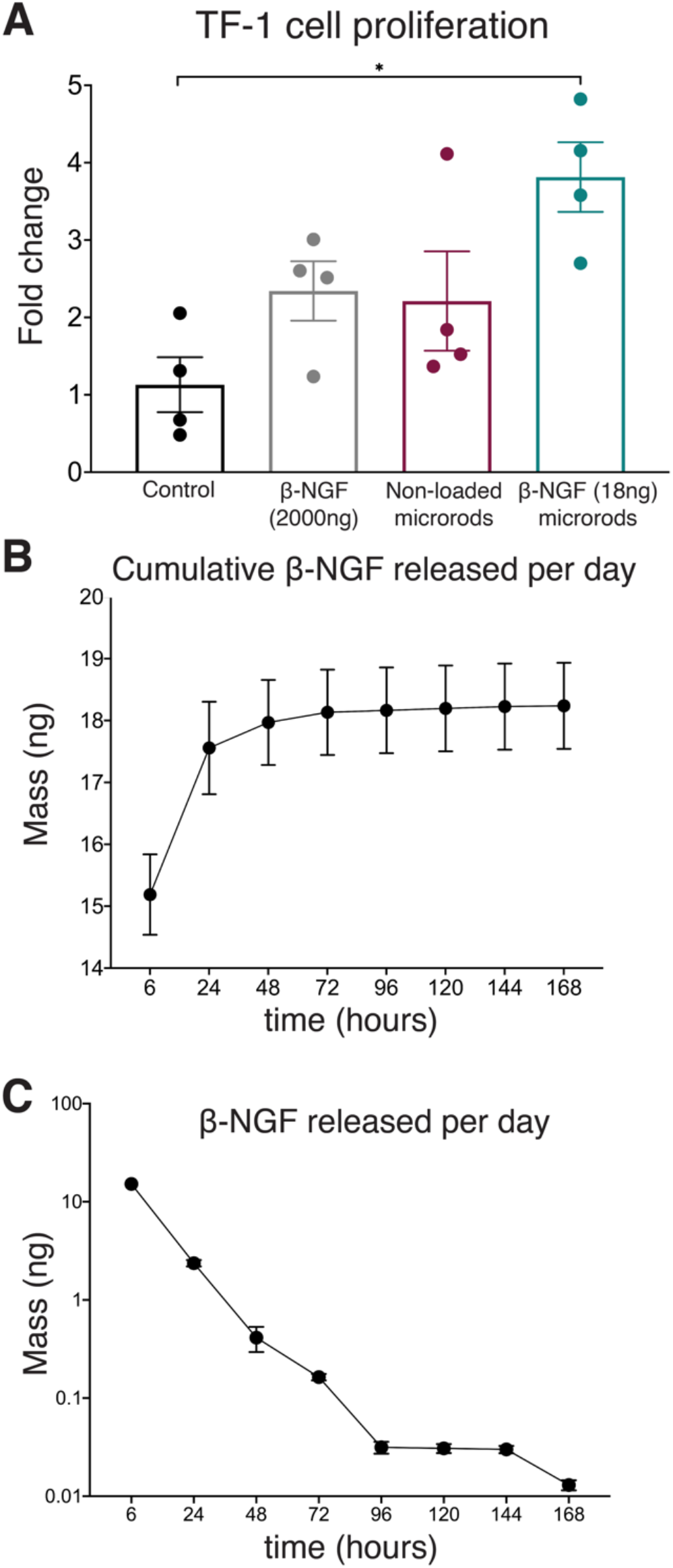
β-NGF loaded onto PEGDMA microrods retain bioactivity. (A) Relative fold change in TF-1 cell proliferation (day 4 vs day 0) for each experimental group. *p<0.05 determined by ANOVA with Tukey’s post hoc test for multiple comparisons (n=4). (B) Cumulative mass (in ng) and (C) daily mass (in ng) of beta NGF released from 90% PEGDMA microrods over a 7-day period shown in hours (n=4). All data shown as means with error bars representing SEM.

### β-NGF loaded PEGDMA microrods promote endochondral bone formation

Our data thus far has demonstrated that PEGDMA microrods can be loaded with β-NGF and β-NGF is bioactive and released over a 7 day period. Given these findings, we hypothesized that sustained release of β-NGF loaded PEGDMA microrods could accelerate endochondral fracture repair. To test this hypothesis, we utilized a murine model of long bone healing wherein closed, mid-shaft fractures were created in the right tibia of adult wild type mice (Supplemental Fig. 2). As previously shown, these non-stabilized fractures incite robust endochondral repair^34,49,51^. In our previous study, we demonstrated that β-NGF was most effective in promoting fracture repair when delivered 7 days post-injury, during the cartilaginous phase of bone healing^34^. Based on this we chose to similarly deliver PEGDMA microrods 7 days post injury using a Hamilton syringe for percutaneous directly to the fracture site. First, we confirmed that the 16,000 microrods suspended in 20 μL saline (lightly stained blue) could effectively be delivered to fracture callus (**Fig. 4A-D)** and remained localized throughout the entirety of the repair period of 14 days **(Fig. 4C-E)** as visualized by Hall-Brunt’s Quadruple (HBQ) staining (cartilage = blue, bone = red). To assess the effectiveness of β-NGF, 16,000 microrods containing 18 ng of β-NGF was injected into the fracture callus. For comparison, additional mice were divided into three experimental (injection) groups: fracture calluses were percutaneously injected with either 20 μL saline, single dose of β-NGF (2,000 ng), or non-loaded PEGDMA microrods. All percutaneous injections were administered 7 days post-fracture and were allowed to heal for 7 days (14 days post-fracture) at which point the calluses were harvested for Micro-Computed tomography (μCT) analysis to quantify the mineralized tissue within the fracture callus and to analyze the bone tissue microarchitecture. By gross examination of the images, the β-NGF microrods group appeared to have a largest most consolidated bony callus compared to all others (**Fig. 5A-D**). Quantification of the bone volume fraction (BFV) confirmed the highest BVF in the β-NGF microrods group with a significantly higher BVF (∼52 % increase) compared to saline controls (**Fig. 5E**). Additionally, the fractures treated with β-NGF microrods resulted in more mature fracture calluses. β-NGF microrods treatment significantly increased trabecular bifurcations (TB, ∼95 % increase) and bone mineral density (BMD, ∼34 % increase) compared to saline controls (**Fig. 5F-G**). Interestingly, the soluble β-NGF did not improve bone formation to the same extent as β-NGF microrods and was not statistically different from the saline controls. Although not statistically different, the non-loaded microrods exhibit higher amounts of BVF, TB, and BMD when compared to the soluble β-NGF and saline treated groups.

**Figure 4.**
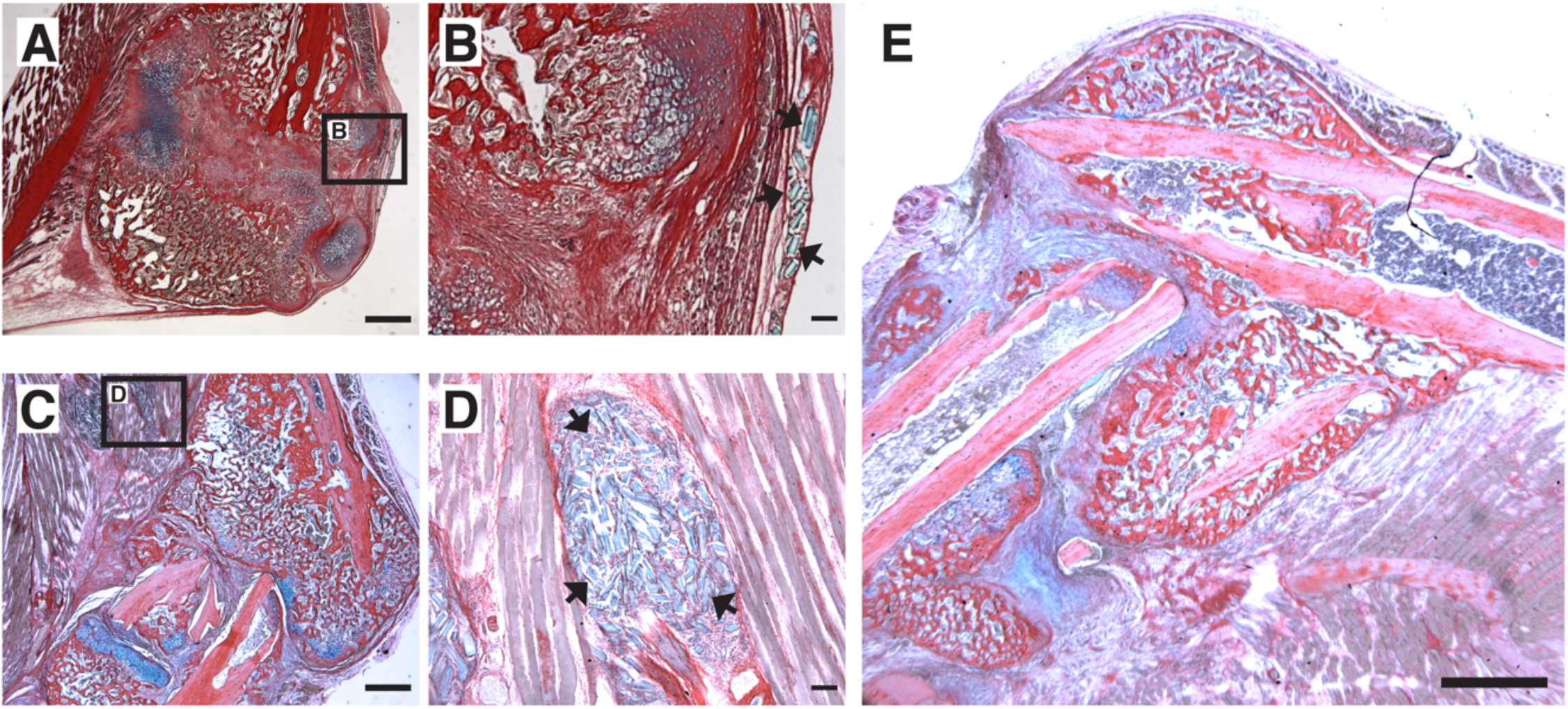
Localization of PEGDMA microrods within tibial fracture calluses. Representative micrographs of (A) low and (B) high magnification of HBQ-stained fracture calluses 5 days post-microrod injection. Representative micrographs of (C) low and (D) high magnification of fracture calluses 7 days post-microrod injection. Arrows indicate PEGDMA microrods within calluses. (A,C) Scale bars = 1mm, (B,D) Scale bars = 100μm. (E) Representative micrograph of fracture callus 14 days post-injection, scale bar = 1mm.

**Figure 5.**
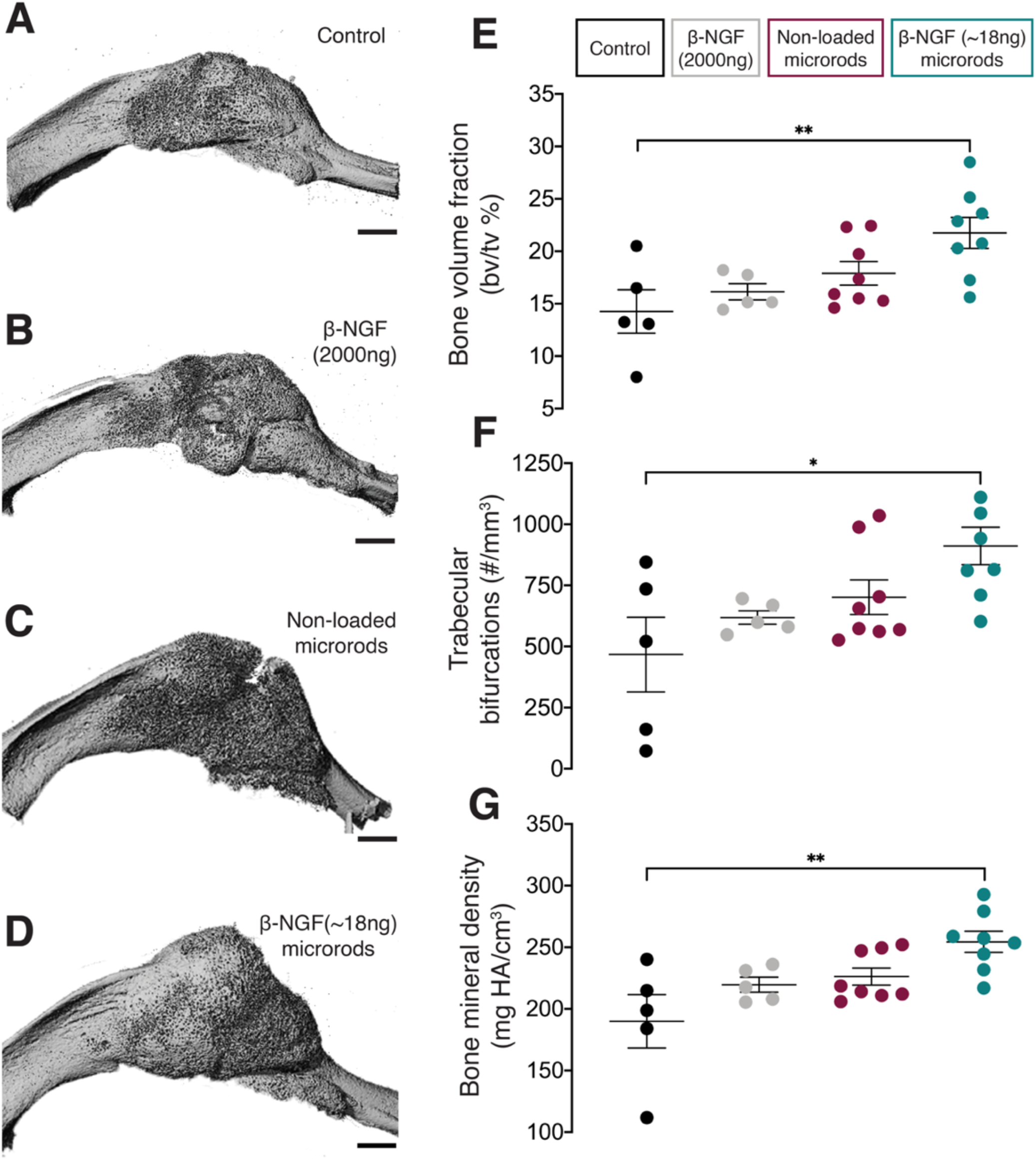
MicroCT analysis of newly formed bone within fracture calluses. Representative three-dimensional images of tibial fracture calluses from mice treated 14 days post-fracture with (A) saline as controls (B) single dose of β-NGF (2000ng) (C) non-loaded PEGDMA microrods and (D) PEGDMA microrods loaded with β-NGF (18ng). Scale bars = 1mm. Quantification of (E) bone volume fraction (F) trabecular connective density and (G) bone mineral density. Error bars represent SEM, *p<0.05, **p<0.01 determined by ANOVA with Tukey’s post hoc test for multiple comparisons.

### β-NGF loaded PEGDMA microrods reduce cartilaginous tissue volume in the fracture callus

To understand the differences noted by μCT at a more detailed tissue level, we utilized quantitative histology to differentiate the cartilage and bone fractions within the fracture callus. Histological images of tibia sections harvested 14 days post-fracture stained with HBQ (cartilage = blue, bone = red) visually indicate that β-NGF microrods have the highest amount of bone (**Fig. 6A-D**). Saline treated fracture calluses had the high quantities of cartilage as percent composition of the callus (32 ± 2 %) with the least bone volume as percent composition (67 ±2 %) compared to other treatment groups indicating the least advanced healing (**Fig. 6A-F**). Higher magnification images verify large proportions of chondrocytes in the fracture callus which suggest the fracture is only nearing the cartilage to bone transition phase in endochondral repair (**Fig. 6A**). Near identical results were observed for the empty microrod treated fracture calluses with cartilage and bone volume at 32 ± 2.7 % and 68 ± 2.7 %, respectively (**Fig. 6C and 6E-F**). The soluble β-NGF treated fracture calluses resulted in slightly elevated levels of bone (71 ± 3.2 %) and lowered cartilage volumes (29 ± 3.2 %) relative to the empty microrods and untreated controls, but this effect was not statistically different. β-NGF loaded microrods were the only treatment group to significantly change the fracture callus composition producing robust bone formation (79 ± 3 %) (**Fig. 6D and 6F**). β-NGF microrod treated samples also show a significant visual reduction in cartilage and statistically different cartilage volume (21 ± 3 %) compared to saline controls (**Fig. 6D and 6E**).

**Figure 6.**
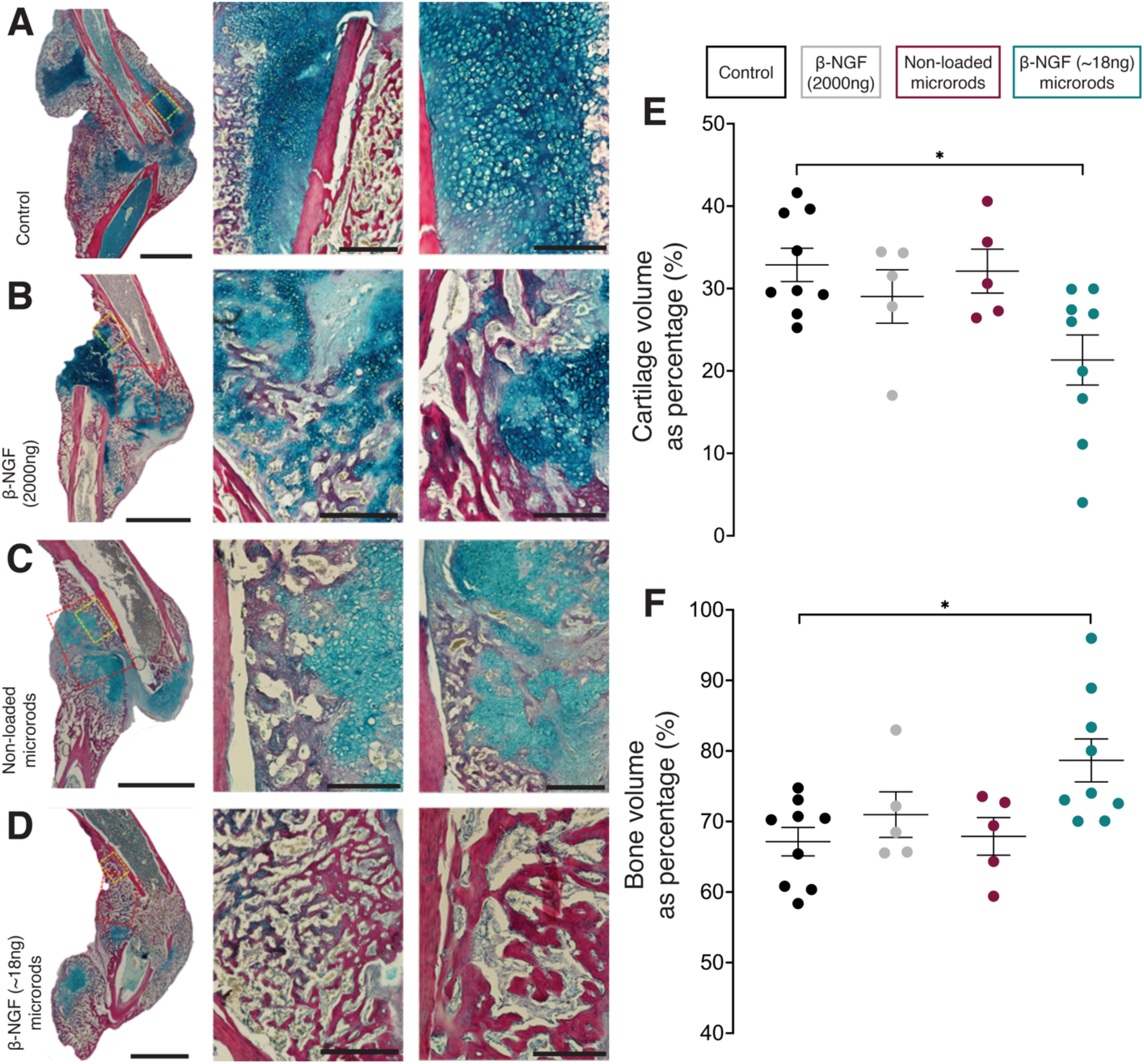
Single injections of PEGDMA microrods loaded with β-NGF promote endochondral bone repair. Representative micrographs of HBQ-stained fracture calluses from mice 14 days post-fracture treated with (A) saline as controls (B) single dose of β-NGF (2000ng) (C) non-loaded PEGDMA microrods and (D) PEGDMA microrods loaded with β-NGF (18ng). Left column scale bars = 2mm, middle and right column scale bars = 500 μm. Quantification of (E) cartilage volume and (F) bone volume, both given as percent composition of fracture callus. Error bars represent SEM, *p<0.05 determined by ANOVA with Tukey’s post hoc test for multiple comparisons.

## Discussion

Recent work in the field supports that NGF signaling plays a significant role in both intramembranous and endochondral ossification. The pioneering mechanistic studies first describe a critical role of NGF/TrkA signaling during skeletal development and adaptation to mechanical load^32,33^. Subsequently our group delineated the spatiotemporal expression of TrkA during long bone fracture repair and identified the cartilaginous phase of healing as a key timepoint for exogenous β-NGF injections by promoting pathways associated with endochondral ossification^34^. Most recently, NGF/TrkA signaling was also shown to be acutely upregulated following stress fracture, triggering reinnervation, vascularization, and osteoblastic activity^31^. Stress fractures are a specific and unique subclass of fracture repair that heal through intramembranous bone formation. Together these studies suggest that NGF could be used as an osteobiologic drug to promote long bone fracture repair by synergistically promoting intramembranous and endochondral bone repair.

This study aimed to build on our previous work by developing a clinically relevant drug delivery system for sustained β-NGF delivery using PEG-based hydrogel microparticles. In our previous study, we locally injected 500 ng of recombinant β-NGF once daily for 3 days to produce a quantifiable acceleration of fracture healing. However, β-NGF is a known hyperalgesic and preclinical studies utilizing murine models have demonstrated that hyperalgesia in mice can be experienced with daily injections of 100 ng and above^52,53^. Thus, our goal is to balance the trophic benefit of β-NGF therapy while minimizing its hyperalgesic effects by providing sustained drug release at a dosing below this threshold. To avoid repeated doses, we aimed to use PEGDMA microrods as a clinically relevant drug delivery platform. The majority of PEGDMA microparticle delivery platforms use spherical particles for bone repair applications^54–56^. Uniquely, in our study we demonstrate the use of high aspect ratio microrods for fracture repair given that high aspect ratio particles have higher residence time and tend to evade phagocytosis or cellular internalization^43,44^. To our knowledge, this is the first study to use PEGDMA microrods for bone fracture repair.

By employing photolithography, PEGDMA microrods can be produced in a high throughput fashion. Moreover, we were able to increase β-NGF loading onto PEGDMA microrods using 90 % (v/v) PEGDMA macromer. Tuning the cross-linking densities can alter the polymer mesh size within the hydrogel and subsequent drug loading and release^36,39^. Higher concentrations of PEG have shown greater loading capacities as seen in our system^55^. Interestingly, striking visual differences in loading are apparent between lower and higher molecular weight molecules such as DAPI and FITC-BSA. DAPI stains the PEGDMA microrods uniformly versus the FITC-BSA that can mostly be visualized on the surface of the PEGDMA microrods immediately after loading. Given that DAPI has a molecular weight of 0.277 kDa compared to the 67 kDa size of FITC-BSA, the micrographs confirmed that molecules of smaller size can more readily diffuse across or into the PEGDMA polymer mesh network.

We next wanted to demonstrate that the loaded β-NGF retained its bioactivity with an *in vitro* proliferation assay using the TrkA expressing TF-1 cell line^46^. Importantly, this assay was performed with 16,000 PEGDMA microrods, versus 100,000 used for the loading assay. This is because only 16,000 could effectively be aspirated in a 20 μL syringe used for the *in vivo* experiments. We calculated that approximately 30-40 % of total protein loaded in 100,000 microrods was about 1-2 mg. Therefore, we used the highest calculation of 2 μg (2,000 ng) to be loaded into the 16,000 microrods and set that as our soluble β-NGF amount for all experiments in parallel. However, the highly specific ELISA assay measured only ∼18 ng of β-NGF in 16,000 PEGDMA microrods. This finding suggests that some of the β-NGF proteins may lose their native molecular arrangement during loading or elution, thus reducing bioactivity. Presumably, the discrepancy between the μBCA assay and the ELISA calculations may be attributed to the disruption of β-NGF’s non-covalent homodimer confirmation or the non-specificity of the μBCA assay. Nonetheless, 16,000 PEGDMA microrods containing 18 ng of bioactive β-NGF’s had a potent effect on TF-1 cell proliferation. This potent effect is likely driven by the sustained release of β-NGF over the 96 hour experimental period. We also observed a nominal increase in proliferation of cells cultured with non-loaded PEGDMA microrods. Although the degradation products of the PEGDMA microrods were not evaluated in this study, PEG at low concentrations have previously been shown to slightly elevate cell proliferation which could be contributing to TF-1 cell proliferation in this study^57^.

Next, we examined PEGDMA microrod localization during endochondral fracture repair in a murine fracture model and were able to histologically localize the PEGDMA microrods at both 5 and 7 days post-injection. However, the PEGDMA microrods were no longer visible after 14 days suggesting that the PEGDMA microrods are perhaps physically degraded overtime due to the dynamic mechanical microenvironment within the fracture callus^58^. Additionally, the fractured leg experiences regular loading due to free ambulation of the animals, which could also contribute to mechanical degradation of the PEGDMA microrods. Degradation products of the PEGDMA microrods are not of concern *in vivo* based on established biocompatibility and non-cytotoxicity^38,40,41^. Importantly, efficacy studies confirm that the PEGDMA microrods were localized in the fracture callus long enough to deliver the therapeutic payload and therefore are suitable for application in fracture callus injections of β-NGF.

Therapeutic efficacy of β-NGF delivery via PEGDMA microrods was validated using non-stabilized tibial fracture in mice followed by μCT analysis and quantitative histomorphometry. β-NGF loaded microrods enhanced endochondral fracture repair as evidenced by the reduction in cartilage volume and statistically significant increases in bone volume fraction (BVF), trabecular bifurcations (TB), and bone mineral density (BMD). Notably, the woven-like bone morphology and minimal hypertrophic chondrocyte cells within the fracture callus further indicates a quicker transition into the bony callus formation after injection with sustained release β-NGF loaded PEGDMA microrods, supporting our hypothesis. We ascertain that this is likely due to the sustained release of β-NGF from the PEGDMA microrods versus a large bolus dose from free/injected β-NGF. Large bolus doses are at risk of off target effects and toxicity^12,35,39^. Thus, sustained release from hydrogels provides a more optimal strategy primarily from a pharmacokinetic approach by slowly delivering drug and maintaining a high local concentration of drug in the target tissue^36,59^.

Another key aspect that could be contributing to the enhanced endochondral bone repair is the likelihood of an increased half-life of β-NGF within the PEGDMA microrods. Hydrogel delivery systems have been described in the literature to improve the half-life of drugs and biologics^39,60,61^. Specifically, β-NGF has an extremely fast distribution half-life of 5.4 minutes and an elimination half-life of 2.3 hours.^62^ Thus, we speculate that the half-life of the successfully loaded β-NGF is extended by encapsulation into the PEGDMA microrods. One important distinction to note is the large difference in β-NGF used in the experimental groups; the soluble β-NGF dose (2,000 ng) was ∼111-fold larger than the β-NGF microrods (18 ng). However, despite the markedly lower dose of β-NGF, the loaded PEGDMA microrods significantly promoted endochondral bone repair compared to the saline treated mice. In addition to potentially extending the half-life of β-NGF, the high localization of β-NGF provided by the PEGDMA microrods may synergistically have contributed to the robust endochondral fracture response seen in mice treated with β-NGF loaded microrods.

## Conclusions

We have demonstrated the feasibility of using PEGDMA microrods for β-NGF sustained delivery and its therapeutic efficacy in a preclinical murine fracture model. We build on our previous data that showed repeated injections of β-NGF during fracture repair promotes endochondral fracture repair. Although that study used multiple injections, this study suggests that lower doses (18 ng) may in fact be more beneficial than larger doses when β-NGF is delivered locally in a sustained manner and encapsulated in a delivery system that may increase the biologics half-life. This dosing regimen of 18 ng is clinically relevant given that NGF has known hyperalgesic effects and thus limiting the total amount of NGF could potentially reduce the pain experienced by subjects. From a therapeutic standpoint, reducing the pain and number of injections is critical given that increased stress can lead to delayed fracture healing^67^. To deliver a sustained dose of β-NGF we utilized PEGDMA hydrogels, as PEG and PEG-conjugates have been FDA approved in the past for several products from tissue adhesives to particle based drug and gene delivery systems^68–70^. To that end, the use of injectable β-NGF loaded PEGDMA microrods validates a novel and translational therapeutic approach for improving bone fracture repair.

## Acknowledgements

The study was supported by National Institutes of Health through NIGMS (R25-GM056847 to K.O.R.), NIDCR (F31-DE028485 to K.O.R., F30-DE031158 to D.L.C.), NIAMS (R01-AR077761 to C.S.B, T.M., and T.A.D.) and the UCSF Academic Senate Committee on Research to C.S.B. and T.A.D. The content is solely the responsibility of the authors and does not necessarily represent the official views of the National Institutes of Health. We would like to thank Nick Szeto and Dr. Wenhan Chang at UCSF’s Core Center for Musculoskeletal Biology & Medicine for μCT analysis.

## Author Contributions

K.O.R, C.S.B., and T.A.D. designed the experiments. K.O.R, B.N.K., and A.N.K. were involved with microfabrication. K.O.R., D.L.C., and A.N.K., performed experiments. K.O.R., D.L.C, V.D., and K.M.O. analyzed the data. K.O.R., D.L.C, C.S.B., and T.A.D. interpreted the results and wrote the manuscript. K.O.R. and D.L.C contributed equally to the manuscript. K.O.R., D.L.C., T.M, C.S.B., and T.A.D. received funding.

