## Supplemental Figures 1-4 for "Localized delivery of β-NGF via injectable microrods accelerates endochondral fracture repair"

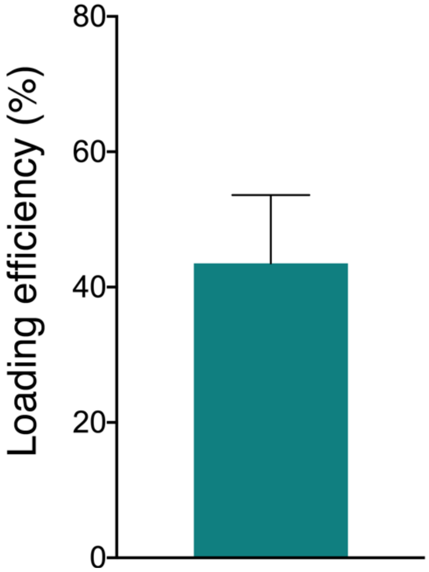


Supplemental Figure 1. Loading efficiency of β-NGF onto 90% PEGDMA (v/v) microrods, measured by microBCA (n=3). Data shown as mean with error bar representing SEM.

A

B

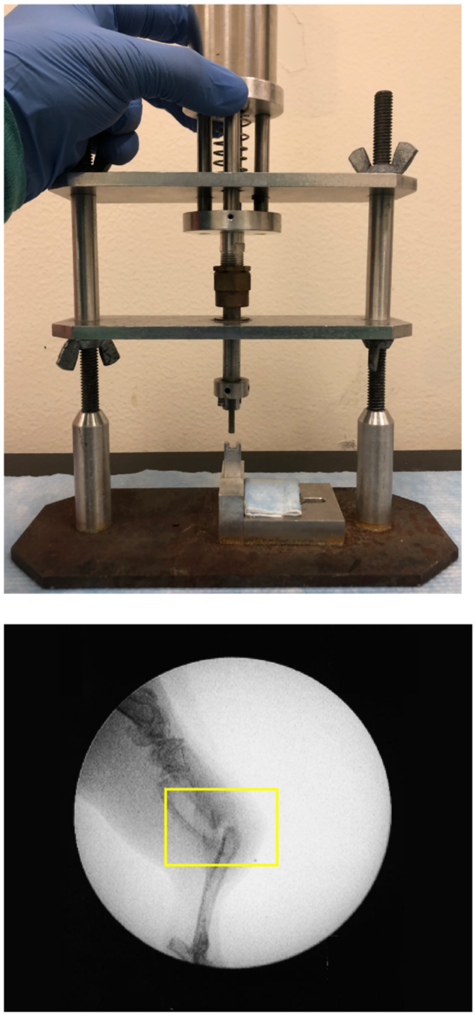

Supplemental Figure 2. (A) Three-point fracture device used to create closed non-stabilized fractures on mouse tibia. (B) Gross fluoroscope image of the entire tibia with yellow frame indicating the mid diaphyseal bone fracture.

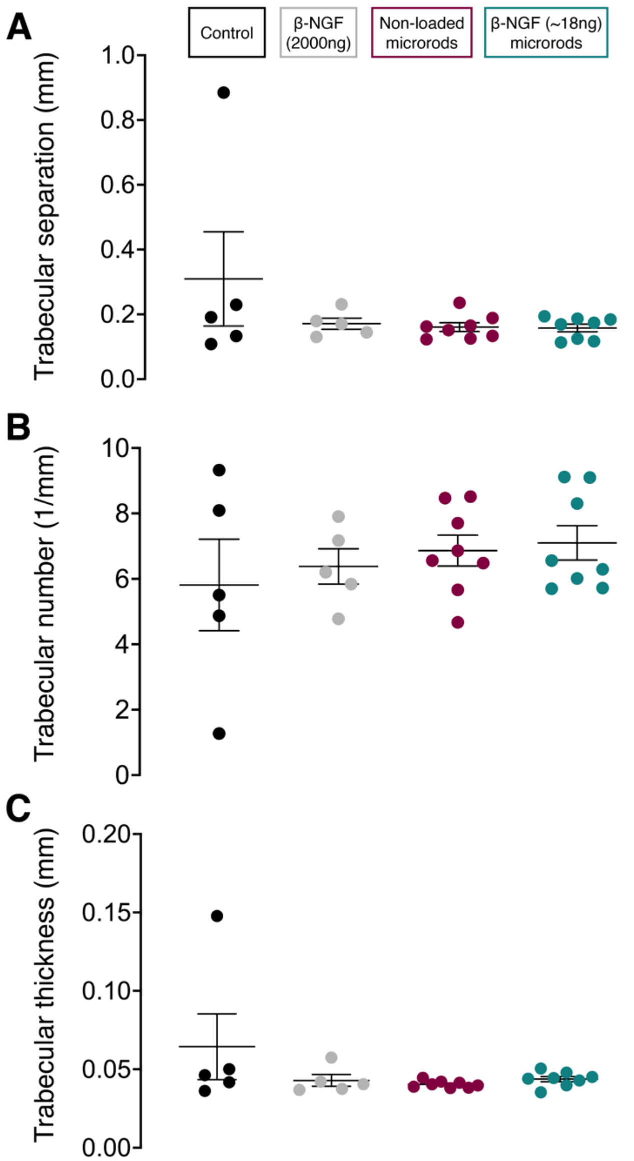


Supplemental Figure 3. MicroCT analysis of trabecular bone within fracture callus. Quantification of (A) trabecular separation (B) trabecular number, and (C) trabecular thickness in mice treated with saline (as control), single dose of β-NGF (2000ng), non-loaded PEGDMA microrods and, PEGDMA microrods loaded with β-NGF (18ng). Error bars represent SEM, non-significance determined by ANOVA with Tukey's post hoc test for multiple comparisons

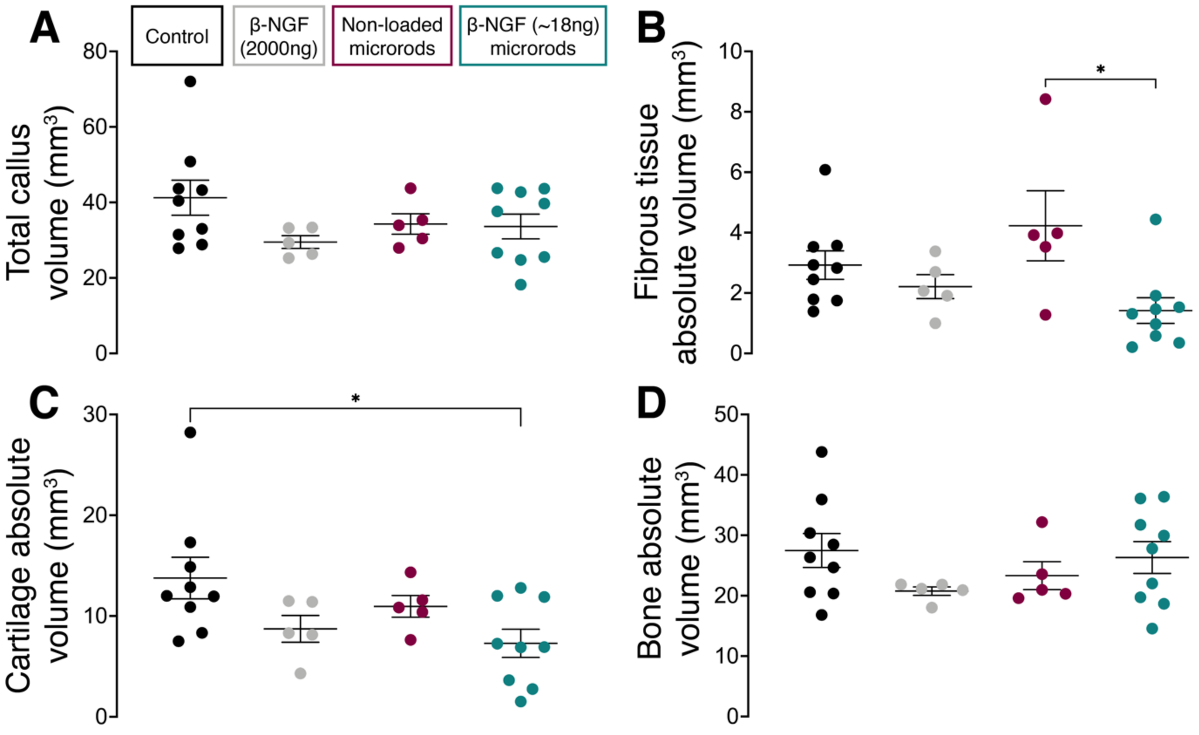


Supplemental Figure 4. Histomorphometric analysis of fracture calluses. Quantification of (A) total callus volume (B) fibrous tissue absolute volume (C) cartilage absolute volume and (D) bone absolute volume in mice treated with saline (as control), single dose of β-NGF (2000ng), non-loaded PEGDMA microrods and, PEGDMA microrods loaded with β-NGF (18ng). Error bars represent SEM, *$p<0.05$ determined by ANOVA with Tukey's post hoc test for multiple comparisons.
